# *Mycobacterium tuberculosis* inhibits autocrine type I interferon signaling to increase intracellular survival

**DOI:** 10.1101/425116

**Authors:** Dallas A. Banks, Sarah E. Ahlbrand, V. Keith Hughitt, Swati Shah, Stefanie N. Vogel, Najib M. El-Sayed, Volker Briken

**Affiliations:** Department of Cell Biology and Molecular Genetics, University of Maryland, College Park, Maryland, USA; Center for Bioinformatics and Computational Biology, University of Maryland, College Park, Maryland, USA; Department of Microbiology and Immunology, University of Maryland School of Medicine, Baltimore, Maryland, USA

## Abstract

The type I interferons (IFN-**α** and -**β**) are important for host defense against viral infections. In contrast, their role in defense against non-viral pathogens is more ambiguous. Here we report that IFN-β-signaling in macrophages has protective capacity against *Mycobacterium tuberculosis* (Mtb) via the increased production of nitric oxide. Furthermore, Mtb is able to inhibit IFN-α/β-receptor-mediated cell signaling and the transcription of 309 IFN-**β** stimulated genes which includes genes associated with innate host cell defense. The molecular mechanism of inhibition by Mtb involves reduced phosphorylation of the IFNAR-associated protein kinases JAK1 and TYK2 leading to reduced phosphorylation of the downstream targets STAT1 and STAT2. Overall, our study supports the novel concept that Mtb evolved to inhibit autocrine type I IFN signaling in order to evade host defense mechanisms.

## Introduction

Type I interferons (IFNs) are innate cytokines that are best known for their ability to induce an anti-viral state in cells (1, 2). Upon binding to their shared receptor, type I IFN receptor (IFNAR), a heterodimer composed of IFNAR1 and IFNAR2 transmembrane proteins, the receptor-associated tyrosine kinases JAK1 and TYK2 are activated, this leads to the phosphorylation and activation of STAT1 and STAT2. Activated STAT1 can homodimerize, translocated to the nucleus and bind to IFN-γ-activated sites (GAS) to promote gene transcription of IFN stimulated genes (ISGs). Alternatively, STAT1 will associate with STAT2 and IRF-9 to form the transcription factor ISGF3 which then translocates to the nucleus to bind to IFN-stimulated response elements (ISRE) of ISG and induce their expression (3, 4).

While type I IFNs clearly have a protective function during viral infection, the role of these cytokines during bacterial or protozoan infections is more ambiguous (2, 4-6). IFN-**β** is detrimental to the host during *Mycobacterium tuberculosis* (Mtb) infections. (7-16) Despite the various outcomes of the type I IFN response to infection it is well documented that many intracellular, non-viral pathogens elicit a host response that leads to the increase in IFN-**β** production (2, 4, 5). Multiple cell-surface (Toll-like receptors) and intracellular (e.g., retinoic acid inducible gene I) receptors recognize microbial products and initiate signaling pathways that activate IRF3, IRF7 or AP1 to induce transcription of type I IFN genes (2, 4, 5). In particular, Mtb gains access to the host cell cytosol via their ESX-1 type VII secretion system, where secreted bacterial DNA (eDNA) binds to the cyclic GMP-AMP (cGAMP) synthase (cGAS) that subsequently activates the STING/TBK1/IRF3 pathway leading to the increased transcription of type I IFNs genes (17-21). The secretion of bacterial c-di-AMP can also mediate the cGAS-independent activation of the STING pathway (22, 23). Finally, Mtb can induce IFN-**β** production through mitochondrial stress and subsequent release of mitochondrial DNA (mtDNA) which activates the STING pathway (24).

The potential of non-viral pathogens to inhibit cell signaling via the IFNAR has not been studied in great detail. One reason for this is probably that the infected host cell detects the pathogen and responds by increased synthesis of IFN-**β** which confounds the analysis. In order to overcome this problem, we used bone marrow-derived macrophages (BMDM) from *Ifn**β***^-/-^ knock-out mice and investigated the effect of IFN-**β** on survival of Mtb and the capacity of Mtb to inhibit IFNAR-mediated cell signaling.

## Materials and Methods

### Cell Culture and Mice

*Ifn-**β***^-/-^ mice were originally obtained by Dr. E. Fish (University of Toronto) and are described in (25). C57BL/6J and *Nos2^-/-^* mice were obtained from The Jackson Laboratory. All animal studies were approved by the IACUC and were conducted in accordance with the National Institutes of Health. Bone marrow-derived macrophages (BMDMs) were prepared from bone marrow cells flushed from the femurs and tibia of mice that were cultured in DMEM supplemented with 10% heat-inactivated FCS, 1% penicillin/streptomycin, and either 20% L929 supernatant for BMDMs during a period of 6 days prior to infection. The Raw264.7-derived, *Irf-3* deficient and IFNAR-signaling reporter cell line (RAW-Lucia™ ISG-KO-IRF3) is commercially available, and measurement of reporter activity was performed according to manufacturer’s protocol (Invivogen).

### Ethics statement

All animals were handled in accordance with the NIH guidelines for housing and care of laboratory animals and the studies were approved by the Institutional Animal Care and Use Committee at the University of Maryland (RJAN1702).

### Bacteria

*M. smegmatis* (mc^2^155), *M. bovis* BCG-Pasteur and *M. tuberculosis* H37Rv (ATCC 25618) strains were obtained from Dr. W. R. Jacobs Jr. (AECOM). *M. kansasii* strain Hauduroy (ATCC 12478) was obtained from ATCC. Bacterial strains were grown in 7H9 media supplemented with 10% ADC, 0.5% glycerol and 0.05% Tween 80. Hygromycin (50 μg/ml) and kanamycin (40 μg/ml) were added to the mutant and complemented strain cultures, respectively.

### *Mycobacterium tuberculosis* (Mtb) *ex vivo* Infection

Bacterial infections of BMDMs were performed as previously described (26). After infection cells were incubated in media containing 100 μg/mL gentamicin in the absence or presence of IFN-**β** or IFN-**γ** (Peprotech). For assays using the RAW-Lucia™ ISG-KO-IRF3 reporter cell line, cells were infected at MOI 10 and stimulated with the indicated amount of IFN-**β**. For measurement of bacterial survival in macrophages, a total of 0.5 million *Ifn-**β***^-/-^, *Nos2*^-/-^, C57BL/6J BMDMs were seeded in 24-well plates and infected with Mtb H37Rv at MOI of 3. Selected BMDMs were then treated with 1000 pg/mL IFN-**β** twice a day for a total of four days. Cells were lysed at indicated timepoints with 0.1% Triton X-100 in PBS, and serial dilutions were plated on 7H11 agar plates (Difco). Colony forming units (CFUs) were counted after 15-20 days of incubation at 37°C.

### Transwell infections

6-well transwells with a 0.4µM membrane (Corning) were allowed to equilibrate in medium overnight before seeding cells. 3×10^6 *Ifn**β**-/-* BMDMs were seeded into the upper transwell and 3×10^6 RAW-Lucia™ ISG-KO-IRF3 cells were seeded in the lower transwell. Infections were performed as described earlier in the upper transwell. Selected conditions were then treated with 200pg/mL IFN-**β** in the upper and lower transwell for the indicated timepoints.

### Western blot analysis

Whole cell lysates were obtained by lysing cells with RIPA buffer containing protease and phosphatase inhibitor cocktails (Roche). Protein concentration was measured using the Piece BCA protein assay kit (Thermo Scientific) and proteins were subjected to SDS-PAGE followed by immunoblotting as described (26). Antibodies were detected binding using SuperSignal West Femto chemiluminescent substrate (Thermo Fisher; 34095) and images were acquired using the LAS-300 imaging system (Fuji). All Western blots were performed at least 3 times and the image of one representative result is shown. ImageJ software (NIH) was used for densitometry quantification as described in figure legends for each Western blot.

### Flow Cytometry

After infection, BMDMs were blocked with 5% FCS and rat anti-mouse CD16/CD32 Fc Block (BD Biosciences, 553141) for 15 min followed by incubation with either PE-conjugated mouse anti-IFNAR1 (Biolegend, 127311) or PE-conjugated goat anti-IFNAR2 (R&D Systems, FAB1083P) for 30 min on ice. PE-conjugated mouse IgG1 (Biolegend, 400111) and goat IgG1 (R&D Systems, IC108P) were used as isotype controls. Protein levels were quantified by acquiring 25,000 cells using the Accuri C6 flow cytometer and software (BD Biosciences). Histograms were processed using FlowJo software version 10 (BD Biosciences).

### Measurement of Nitric Oxide (NO) production

NO production was quantified in cell culture supernatants by the Griess reagent kit which measure the NO derivate Nitrite (ThermoFisher, G7921) at the indicated timepoints.Absorbance was measured at 548nm using a microplate reader (BioTek). Nitrite concentration was determined using a sodium nitrite standard curve (0-100 µM). Culture supernatants were pooled from three replicate wells per experiment for all infections. All samples were assayed in technical duplicates and three independent experiments were performed.

### RNAseq Library Preparation and Analysis

*Ifn-**β***^-/-^ BMDMs were infected or not as indicated previously with Mtb H37Rv. At 4 hours post infection (hpi) cells were lysed with 1 mL Trizol (Ambion). RNA was extracted with chloroform, precipitated with 100% isopropanol, and washed with 70% ethanol. Purified RNA was treated with Turbo DNAse (Ambion) for 1 hour.

RNAseq libraries were prepared using the Illumina ScriptSeq v2 Library Preparation Kit according to the manufacturer’s protocol. Library quality was assayed by Bioanalyzer (Agilent) and quantified by qPCR (KAPA Biosystems). Sequencing was performed on an Illumina HiSeq 1500 generating 100bp paired-end reads. RNA-Seq read quality was assessed using FastQC (http://www.bioinformatics.babraham.ac.uk/projects/fastqc/) and low-quality base-pairs were removed using Trimmomatic (27). The Ensembl *Mus musculus* GRCm38 reference genome (version 76) was downloaded from the Ensembl website (28) and reads were mapped to the genome using TopHat2 (29). HTSeq (30) was used to quantify expression as the gene level. Count tables were loaded into R/Bioconductor (31). log2-transformed, counts-per-million (CPM) and quantile normalized (31), and variance bias was corrected for using Voom (32). Next, batch adjustment was performed using ComBat (33). We performed Pearson correlation, Euclidean distance and PCA analyses which revealed the presence of a single outlier Mtb sample (HPGL0627), which was removed from subsequent analyses (Table S1). The differential gene expression was measured for each of several contrasts: 1) uninfected (UI) vs. uninfected +IFN-**β** (UI +IFN-**β**) and 2) Mtb + IFN-**β** vs. UI + IFN-**β** (Table S1). In order to determine which IFN-**β**-stimulated genes were specifically deregulated during infection with Mtb, the intersection of the set of genes found to be differentially expressed in both the UI vs. UI + IFN-**β** and Mtb + IFN-**β** vs. UI + IFN-**β** contrasts was taken (Table S1). All raw RNAseq data were submitted to SRA and can found at https://www.ncbi.nlm.nih.gov/sra/SRP130272.

### Cytokine Measurements

*Ifn-**β***^-/-^ BMDMs were infected as described previously and treated with 50 pg/mL IFN-**β**. An additional 50 pg/mL of IFN-**β** was added at 3 hpi, and cell culture supernatants were collected at 6 hpi. Concentrations of selected cytokines were determined using a custom ProcartaPlex magnetic bead-based multiplex assay (Thermo Fisher Scientific) on the Luminex MAGPIX^®^platform according to manufacturer’s instructions. CCL12 and CCL3 protein levels were measured using ELISAs (R&D Systems)

### Statistical Analysis

Statistical analysis was performed on at least three independent experiments using GraphPad Prism 7.0 software and one-way ANOVA with Tukey’s post-test or Student’s t-test, and representative results of triplicate values are shown with standard deviation unless otherwise noted in the legends. The range of p-values is indicated as follows: * 0.01<p<0.05; **0.001<p<0.01, *** 0.0001<p<0.001, and **** p<0.0001.

## Results

### IFN-**β** has anti-microbial activity via induction of Nos2

The analyses of the importance of IFN-**β** for host defense during infection with non-viral pathogens has proven to be complex since during *in vivo* infections host genetic components, tissue environment and actual doses of IFN-**β** all can affect outcomes (2, 4, 6). In order to investigate the importance of IFNAR-signaling we used a reductionist approach by eliminating bystander cell, host tissue effects and the host cell production of IFN-**β** in response to infection by using BMDM from *Ifn-**β***^-/-^ mice (25). We infected *Ifn-**β***^-/-^ BMDMs with *Mycobacterium tuberculosis* (Mtb), in the presence or absence of 1ng/mL IFN-**β**. The bacterial burden was measured every 24 h for a total of 96 h by counting colony forming units (CFUs). During Mtb infection, we found that reduction in bacterial burden in IFN-**β**-treated cells was minimal at 24 hpi and 48 hpi, but steadily increased between 48 hpi and 96 hpi. At 96 hpi, IFN-**β** treatment reduced bacterial burden by ~50% (Figure 1A). We also determined that IFN-**β**-treated, Mtb-infected cells exhibited less necrosis compared to untreated infected cells since an increased release in gentamycin containing medium could have otherwise accounted for the reduction of CFU (Figure 1B). These results demonstrate that IFN-**β** treatment promotes host defense against intracellular microbes during infection of primary macrophages. The direct antimicrobial activity of IFNAR-signaling on Mtb viability that we show here has not been demonstrated before in a system in which the infection itself does not produce IFN-**β**.

**Fig 1.**
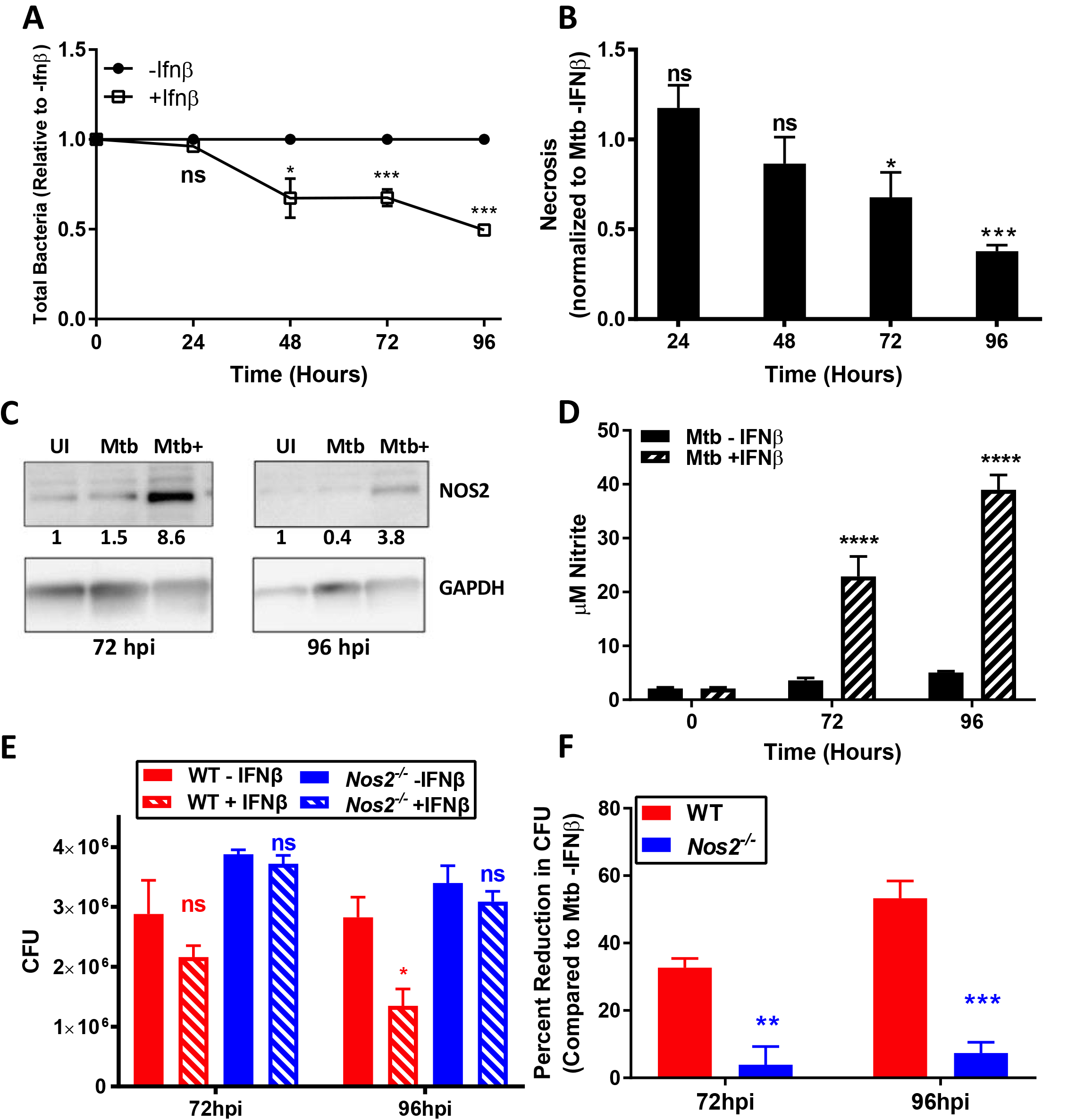
Ifn-**β** treatment has a broad anti-microbial activity. (A) *IFN-**β***^*-/-*^ BMDMs were infected with Mtb H37Rv in the presence or absence of 1000pg/ml IFN-**β**. The number of bacteria by CFUs in treated conditions relative to untreated cells was then calculated. (B) Cell necrosis was assayed at each indicated time point using an adenylate kinase release assay (Toxilight bioassay, Lonza) and is represented as fold change over untreated infected cells. (C) Whole cell lysates were collected at the indicated timepoints and immunoblotted for NOS2. Band densities were normalized to GAPDH. (D) Nitrite levels from culture supernatants were determined using the Griess reagent. (E, F) WT or *Nos2^-/-^* BMDMs were infected with Mtb H37Rv in the presence or absence of 1000pg/ml IFN-**β**. Bacterial burden was determined as in Fig. 1A. All data shown are presented as mean ± S.E.M. of at least three independent experiments.

It was previously shown that type I IFN signaling may lead to the induction of nitric oxide synthase 2 (*Nos2*) gene expression and subsequent nitric oxide (NO) production infected BMDMs (34). In order to investigate whether NO production plays a role in IFN-**β**-mediated clearance *ex vivo*, we determined if NOS2 was expressed at time points correlating with the reduction in bacterial burden during Mtb infection. At 72 hpi, NOS2 was upregulated in Mtb-infected cells, and NOS2 levels were further increased in Mtb-infected cells stimulated with IFN-**β** (Figure 1C). At 96 hpi the NOS2 protein levels were decreased compared to 72 hpi, however there was still a significant upregulation in Mtb-infected cells stimulated with IFN-**β** compared to uninfected cells (Figure 1C). Consequently, we measured NO levels in the different experimental groups in order to assess the effect of IFN-**β** and infection on NO production. In Mtb-infected cells treated with 1ng/ml IFN-**β**, we noticed a sharp increase in nitrite levels which is consistent with the observed increase in NOS2 levels (Figure 1D). We then investigated whether or not IFN-**β** could promote host resistance in *Nos2*^-/-^ BMDMs. We infected wild-type (WT) or *Nos2^-/-^* BMDMs with Mtb in the presence or absence of 1ng/ml IFN-**β**. Infected BMDMs showed a significant decrease in CFUs at 96 hpi in IFN-**β** treated cells (Figure 1E-F). In contrast, we measured no significant decrease in IFN-**β**-treated *Nos2^-/-^* BMDMs at either 72 hpi or 96 hpi (Fig 1E-F). These results demonstrate that IFN-**β** treatment promotes host resistance to Mtb via the production of NO at later time points during *ex vivo* infection.

### Mtb inhibits IFNAR-signaling

Viruses are well known to evade host protective IFN-**β** responses by suppressing signaling via IFNAR (1, 2). Our results on the host protective effect of IFN-**β** on Mtb infection prompted us to investigate the potential of Mtb to inhibit IFNAR-signaling. To this purpose we used a reporter RAW264.7 cell line (Invivogen) which is deficient in *Irf-3* and has an ISG promoter in front of a reporter gene for easy quantification of IFNAR-signaling. We first determined the capacity of these pathogens to induce activation of the reporter in the absence of an external IFN-**β** stimulus. Msm, Mtb H37Rv, Mtb CDC1551 did not induce activation of the ISG reporter in the absence of external stimulation. We also determined that the infection with the different mycobacterial species does not induce the production of IFN-**β** via ELISA since these cells are deficient in IRF-3 (not shown). Next, we added increasing amounts of IFN-**β** to infected or uninfected cells and normalized the response to the response obtained in uninfected cells (Figure 2A). Overall, both virulent Mtb strains consistently showed about 50-60% of inhibition whereas infection with Msm had only a minor effect (10-15%) at higher doses of IFN-**β** (Figure 2A). Notably, this effect was reversed at 1000pg/ml, suggesting there is a limit to the amount of signaling Mtb can inhibit. To our best knowledge the described experiments are the first to demonstrate the capacity of Mtb to inhibit IFNAR-signaling. The lack of investigation into this capacity of the pathogens might be explained by the proposed overall role of type I IFN in exacerbation of disease outcomes (2, 4-6).

**Fig 2.**
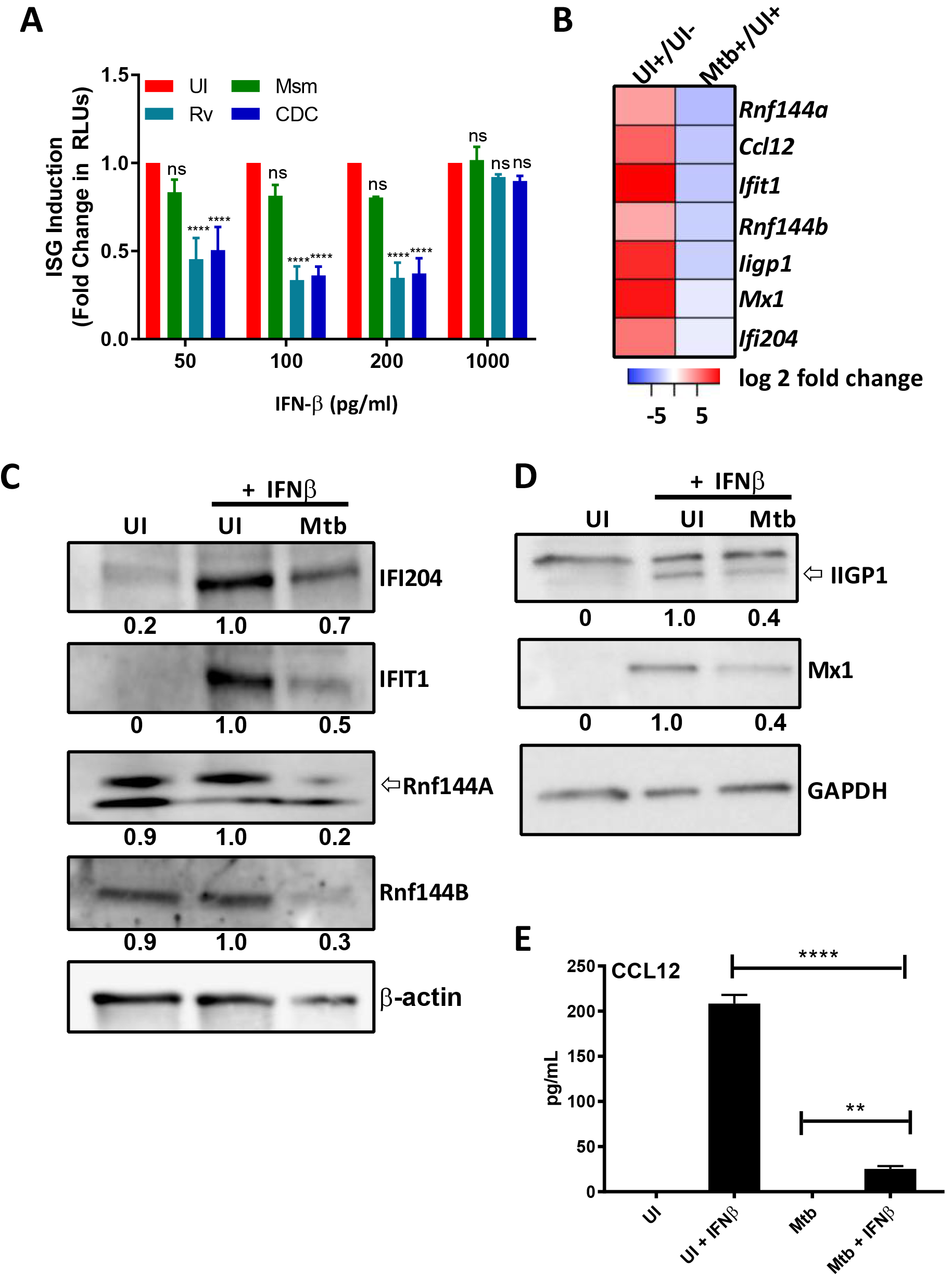
Mtb inhibits type I IFN signaling. (A) IRF-3 deficient RAW264.7 macrophages transfected with an IFN-**β** responsive luciferase gene (Invivogen) were infected and treated with the indicated concentrations of IFN-**β**. After 20hpi the amount of secreted luciferase was quantified. ISG induction is represented as the fold change in RLUs compared to uninfected cells. (B) Heatmap of down-regulated genes selected for follow-up studies. Log-transformed expression ratios for Mtb + IFNβ/UI + IFN-**β** are plotted for each gene. (C-E) *Ifn-*β^*-/-*^ BMDMs were infected with *M. tuberculosis* H37Rv (Mtb) and treated with 50 pg/mL IFN-**β** for 4 hours. (C) Whole cell lysates were collected and immunoblotted for IFI204, IFIT1, Rnf144A, Rnf144B. Band density was normalized to β-actin. (D) Cell lysates were collected and immunoblotted for either MX1, IIGP1, and normalized to GAPDH. (E) Supernatants were analyzed for levels of CCL12 using ELISA. All data shown are presented as mean ± S.E.M. of at least three independent experiments.

### Mtb affects the type I IFN-stimulated gene host transcriptional profile

Signaling via the IFNAR induces the formation the IFN-stimulated gene factor 3 (ISGF3) complex and or STAT1:STAT1 homodimerization which both translocate into the nucleus and bind to ISRE or GAS, respectively, leading to the transcription of hundreds of genes involved in a variety of immunological functions (3). Consequently, we hypothesized that by inhibiting type I IFN signaling, Mtb can manipulate the expression of host genes to promote its intracellular survival. To investigate this, we used RNA sequencing technology (RNAseq) to identify IFN-**β**-regulated genes that are up- or down-regulated during *Mtb* infection in *Ifn-**β***^-/-^-derived BMDMs.We analyzed the following experimental groups: 1. uninfected, untreated (UI), 2. uninfected IFN-**β** treated (UI+) and 3. Mtb H37Rv infected and treated (Mtb+) BMDMs (Table S1). The RNAseq data was reproducible as indicated by principle component analysis (PCA), hierarchal clustering and Pearson correlation (Figure S1). All three analyses depicted a satisfactory degree of clustering between biological replicates with the exception of one outlier (HPGL0627), which was excluded from further analysis (Figure S1 and Table S1). In order to identify IFN-**β**-stimulated genes that exhibited differential expression in Mtb-infected BMDMs, we first characterized all of the IFN-**β**-regulated genes in our experimental system. To that purpose we compared the gene expression of uninfected, untreated (UI) to uninfected, IFN-**β** treated (UI+) conditions. We identified a total of 1144 IFN-**β**-stimulated genes and 956 genes with reduces expression (> 2fold change) (Figure S2A, Table S1). Next, in order to identify all of the genes that are impacted by Mtb-infection we compared gene expression levels between the UI +IFNβ condition and the Mtb H37Rv infected and treated (Mtb+) and found 1296 upregulated and 1294 downregulated (> 2fold change and an adjusted p-value <0.05) genes (Figure S2B, Table S1). Finally, Mtb-infection causes the deregulation of many genes that are not regulated by IFN-**β** but will be included in the UI+ versus Mtb+ contrast. The overlap between these two gene sets was determined to be 309 genes with reduced expression and 170 with increased expression for a total of 479 deregulated genes (Figure S2C, Table S1). To extend the results of our RNAseq analysis, we determined protein expression and secretion levels of selected targets in *Ifn-**β***^-/-^ BMDMs infected with Mtb in the presence or absence of IFN-**β** (Figure 2B). The selection of candidate gene products for follow-up studies was based on magnitude of deregulation and availability of antibodies (see list of 479 genes in Table S1). Rnf144A and Rnf144B proteins were not significantly upregulated upon treatment with IFN-**β** (Figure 2C) which may reflect a low fold-increase at the RNA level as illustrated in the heatmap (Figure 2B), however IFI204, IFIT1, MX1 and IIGP1 were strongly induced after treatment with IFN-**β** (Figure 2C-D). For all targets, we observed some degree of downregulation in IFN-**β**-treated cells infected with Mtb. In addition, we used ELISA to demonstrate the strong inhibition of IFN-**β**-driven secretion of CCL12 by Mtb infection (Figure 2E). In conclusion, these data show a strong correlation of the RNAseq analysis with protein data for the subset of 309 IFN-**β**-regulated genes that are repressed by Mtb infection (Figure 2B and Table S1). We additionally investigated some of the 170 genes that were strongly upregulated in the 479 genes set (Table S1) and characterized several cytokine/chemokine targets using a combination of both multiplexed and regular ELISA and determined that Mtb infection of BMDMs without addition of IFN-**β** upregulated secretion levels of all assayed proteins (Figure S3). For IL-1β, TNFα and CXCL1 there seemed to be additive effected in Mtb +IFN-**β** treated cells when compared to UI +IFN-**β** alone and Mtb -IFN-**β** mediated cytokine induction, whereas there was an antagonistic effect of the Mtb-infection for IL-27, IL-1α, CCL5 and CCL3 suggesting that the induction of these cytokines in Mtb-infected cells is independent of IFNAR-signaling and that their upregulation caused by addition of IFN-**β** can be inhibited by Mtb (Figure S1). Overall, this analysis shows that this subset of 170 of Mtb-deregulated genes after IFN-**β** addition that is upregulated provides less insights because many of the genes seem to be upregulated by Mtb infection alone.

### Mtb inhibits type-I but not type-II IFN-mediated activation of TYK2, JAK1

We discovered that Mtb inhibits signaling via IFNAR and now we investigated further at what level of the signaling cascade the inhibition occurs. Several viral pathogens have evolved mechanisms to evade the IFN-**β** response by promoting the degradation of IFNAR (35, 36). We found that surface expression levels of IFNAR1 and IFNAR2 using flow cytometry remained unchanged by Mtb infection (Figure 3A). After stimulation of IFNAR1/R2 by IFN-**β**, the cytosolic protein tyrosine kinases JAK1 and TYK2 are recruited and phosphorylated. We showed that Mtb-inhibited tyrosine phosphorylation of both TYK2 and JAK1 as early as 20 min post infection (Figure 3B, 3C). Since JAK1 is involved in both type I and type II interferon signaling, we also analyzed phosphorylation levels of JAK1 and JAK2, another IFN-**γ**-activated protein kinase. In IFN-**γ**-treated and Mtb-infected cells there was no significant change in JAK1 or JAK2 phosphorylation levels, showing that the inhibition of JAK1 phosphorylation is specific to type I IFN signaling (Figure 3D, 3E).

**Fig 3.**
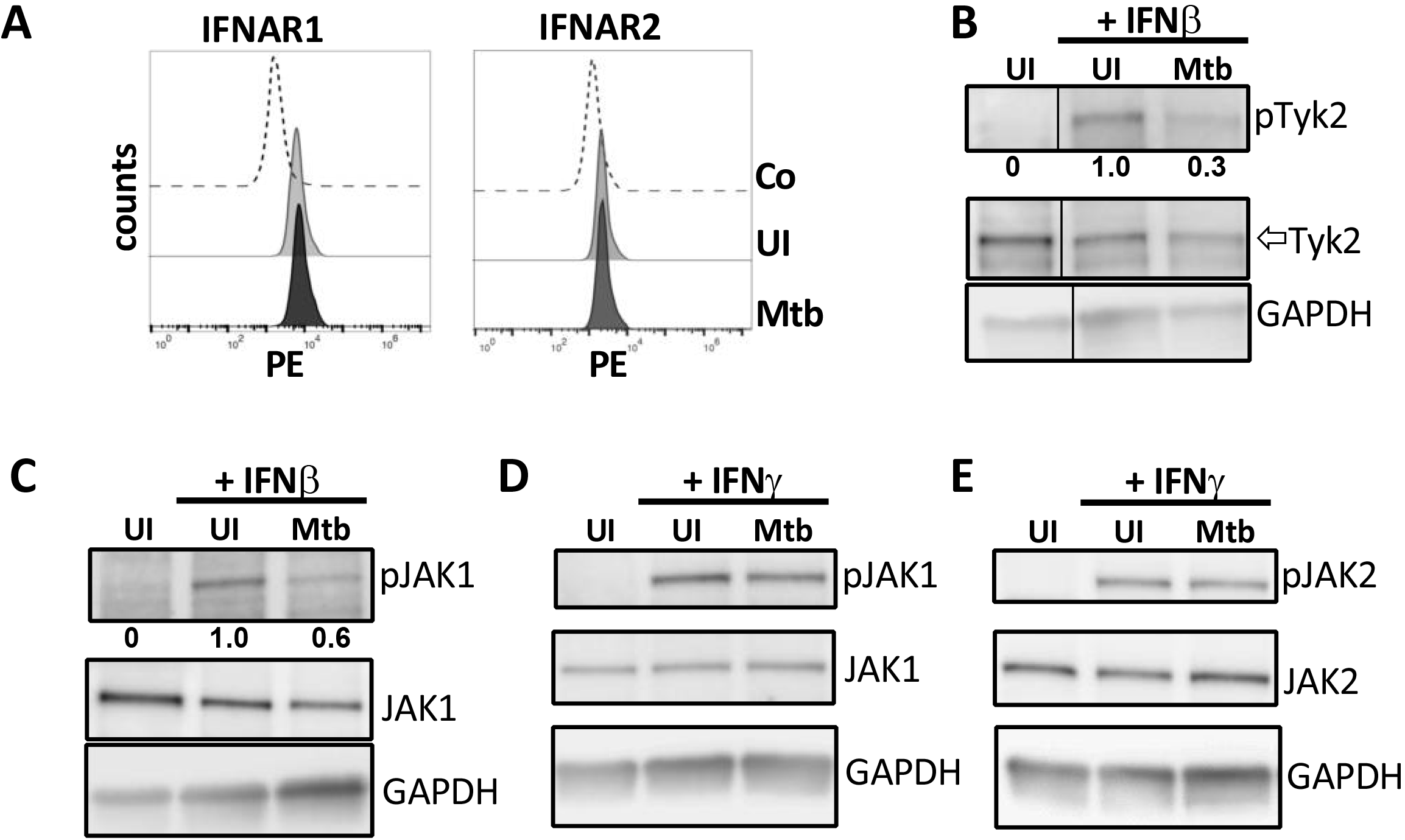
Mtb inhibits type-I but not type-II IFN-mediated activation of TYK2 and JAK1. (A) *Ifn-**β***^*-/-*^ BMDMs were infected with Mtb H37Rv and flow cytometry was conducted at 4hpi to measure surface receptor expression levels of IFNAR1 and IFNAR2. (B and C) *Ifn-**β***^*-/-*^ BMDMs were infected as described in the presence of 300pg/ml IFN-**β**. Cell lysates were collected at 20 min post infection and immunoblotted for pJAK1 (Y1022/1023), total JAK1, pTYK2 (Y1054/1055), and total TYK2. (D and E) *Ifn-**β***^*-/-*^ BMDMs were infected as described in the presence of 300pg/ml IFN-**γ**. Cell lysates were collected at 20 min post infection and immunoblotted for pJAK1 (Y1022/1023), total JAK1, pTYK2 (Y1054/1055), and total TYK2. Densitometry was performed using ImageJ software and phosphorylated protein bands were normalized to total signal for each condition. Data and densities shown represent one representative experiment out of three.

### Mtb inhibits downstream phosphorylation of STAT1 and STAT2

Type I and type II IFNs induce transcription of different subsets of genes but STAT1 phosphorylation occurs in both signaling pathways (3). The canonical signaling pathways are defined as follows: type I IFN induces the heterodimerization of STAT1 and STAT2, while type II IFN (IFN-**γ**) induces homodimerization of STAT1, although type I IFN can also induce STAT1 homodimerization (3). We performed an infection comparing STAT1 tyrosine phosphorylation levels in IFN-**β**-treated *Ifn-**β***^-/-^ BMDMs infected with virulent Mtb strains H37Rv or strain CDC1551 and observed that both could similarly inhibit STAT1 phosphorylation, suggesting that this mechanism of host cell manipulation is shared among virulent Mtb strains (Figure 4A). We also showed that Mtb did not inhibit IFN-**γ**-dependent phosphorylation of STAT1 (Figure 4B) which is consistent with previously published results (37) and suggest a separate molecular pathway engaged by Mtb for the inhibition of IFNAR-signaling.

**Fig 4.**
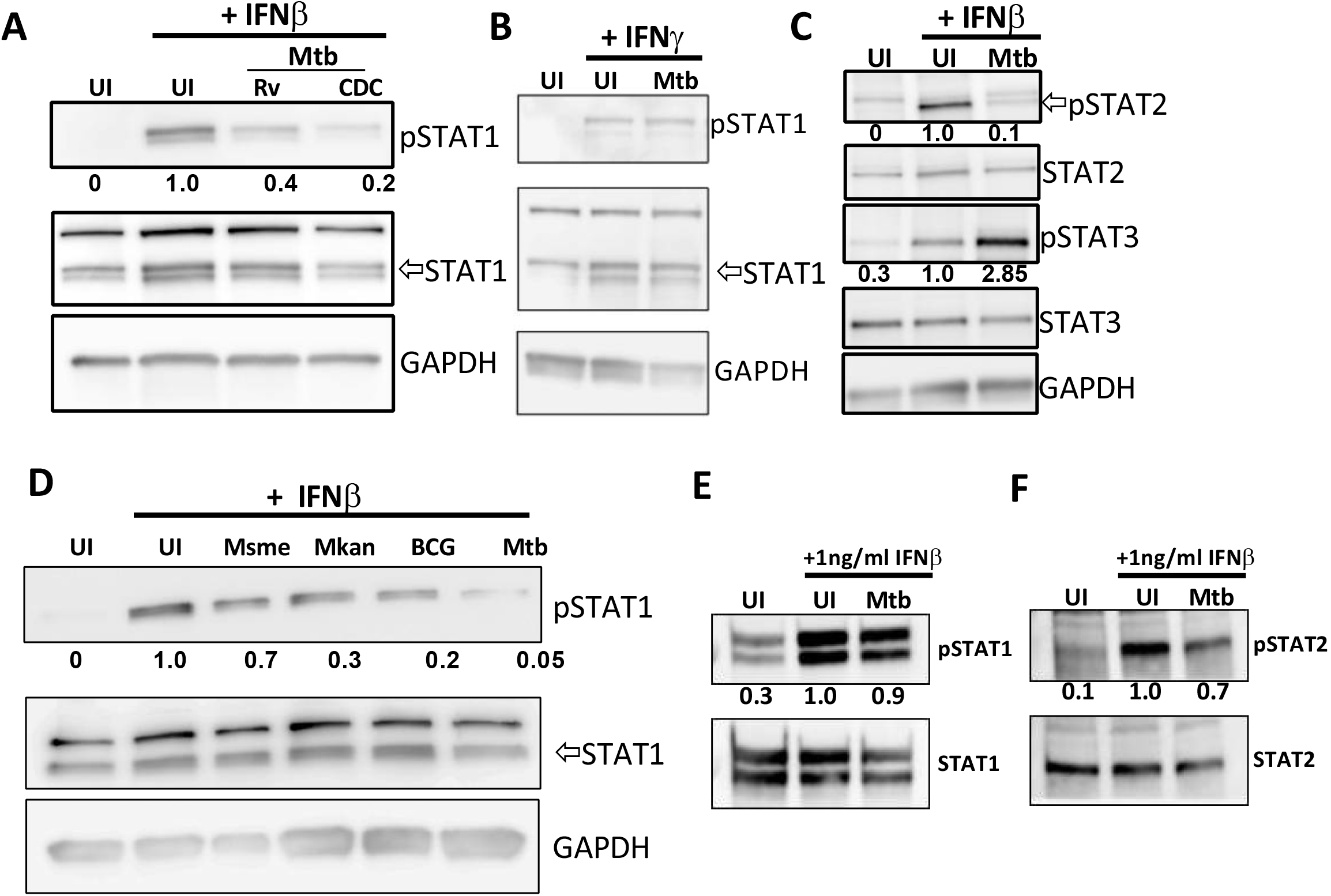
Mtb inhibits phosphorylation of STAT1 and STAT2. (A-B) *Ifn-**β***^*-/-*^ BMDMs were infected with either Mtb strains H37Rv or CDC1551 and treated with 50pg/ml IFN-**β** (A) or 50pg/ml IFN-**γ** (B). Whole cell lysates were collected at 4 hours post infection and immunoblotted for pSTAT1 (Y701) and total STAT1. (C) *Ifn-**β***^*-/-*^ BMDMs were infected with Mtb in the presence or absence of 300pg/ml IFN-**β**. Whole cell lysates were collected at 20 min post infection and immunoblotted for pSTAT2 (Y690), total STAT2,pSTAT3 (Y705), or total STAT3 as indicated. (D) Cells were infected with the indicated *Mycobacteria* strains and treated with 50pg/ml IFN-**β**. Whole cell lysates were collected at 4 hours post infection and immunoblotted for pSTAT1 and total STAT1. (E-F) *Ifn-**β***^*-/-*^ BMDMs were infected with Mtb and treated with 1ng/ml IFN-**β**. Whole cell lysates were collected at 4 hours post infection and immunoblotted for pSTAT1, total STAT1, pSTAT2 or total STAT2 as indicated.

Besides STAT1, other STAT isoforms may be stimulated by type I IFN signaling. STAT2 becomes phosphorylated and heterodimerizes with pSTAT1 and IRF9 to form the ISGF3 complex, which translocates into the nucleus to induce transcription of genes containing interferon stimulated regulatory elements (ISREs) (3). In addition, it has also been shown that STAT3 can be induced by type I IFNs to regulate transcription of different gene subsets (38). We discovered that Mtb also inhibited the tyrosine phosphorylation of STAT2 (Figure 4C). However, Mtb actually induced phosphorylation of STAT3 (Figure 4C). STAT3 phosphorylation can inhibit type I IFN-mediated signaling, although this occurs at the level of nuclear translocation (39, 40). Thus, STAT3 activation is most likely not involved in our observed inhibition of TYK2/JAK1 phosphorylation by Mtb.

The levels of IFN-**β** production by mycobacteria-infected cells vary depending on the mycobacterial species and, in the context of Mtb, the specific strain that is used for the infection (24, 41, 42). We sought to test if the difference in type I IFNs production observed for the different mycobacterial species correlated with their variable capacity to inhibit IFNAR-mediated cell signaling. Here we observed a more modest decrease in the relative STAT1 phosphorylation in cells infected with the vaccine strains *M. bovis* BCG (BCG) and *M. kansasii* but only a minor reduction upon infection with *M. smegmatis* (Fig 4D). Notably, the capacity of Mtb to inhibit STAT1 and STAT2 phosphorylation was also reversed upon addition of a high dose of IFN-**β**, supporting the data using our reporter cell line (Fig 4E-F).

### Mtb infection does not inhibit IFN-**β** signaling in bystander cells

In order to determine whether inhibition of IFN-**β** signaling is specific to Mtb infected cells, and not due to the secretion of a soluble host factor, we sought to determine if there was an effect in bystander cells. To address this, we used a transwell system in which the upper transwell was seeded with *Ifn-**β***^-/-^ BMDMs and the lower transwell was seeded with our ISG reporter cell line (Fig 5A). The upper transwell was then infected or not with Mtb, and both wells were stimulated or not with IFN-**β** as earlier. In this system, a reduction of secreted luciferase from the cells in the lower transwell would suggest that there is indeed a bystander effect. In our hands, we saw equal levels of reporter activity in bystander cells exposed to both uninfected and infected *Ifn-**β***^*-/-*^ BMDMs, suggesting that the inhibition is specific to infected cells (Fig 5B). Western blots of *Ifn-**β***^*-/-*^ BMDM lysates (Fig 5C) and the ISG reporter cell line (Fig 5D) confirm that we do still see inhibition of STAT1 phosphorylation in infected *Ifn-*β^*-/-*^cells, however, STAT1 phosphorylation of bystander cells is not affected by the infection status of the *Ifn-*β^*-/-*^ BMDMs.

**Fig 5.**
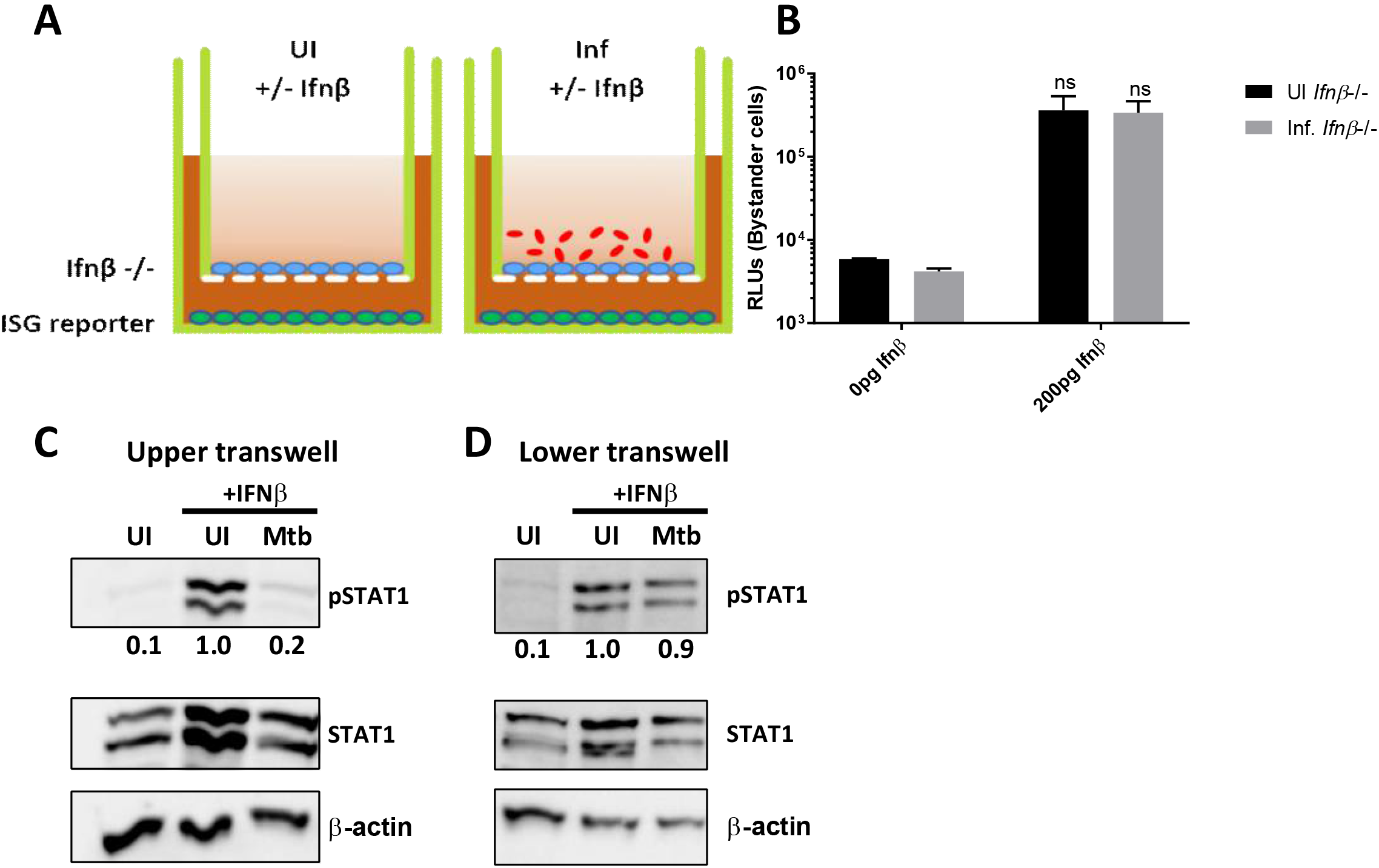
Mtb does not inhibit type I IFN signaling in bystander cells. (A) Schematic of transwell experiments. (B) Upper transwells were infected or not with Mtb, and then the upper and lower transwells were treated with 200pg/ml IFN-**β** for 4 hours. Cells in the lower transwell were lysed by addition of Triton-X-100, and the amount of luciferase was quantified. (C) Whole cell lysates from the upper transwell of *Ifn-**β***^*-/-*^ BMDMs were collected at 4hpi and immunoblotted for pSTAT1, total STAT1 or actin as indicated. (D) Whole cell lysates from the lower transwell were collected at 4hpi and immunoblotted for pSTAT1, total STAT1, or actin as indicated. UI or Mtb refers to the infection condition of the upper transwell.

## Discussion

While type I IFN are largely considered to be beneficial in the context of viral infections, their role during bacterial infections is not completely understood and may vary depending on the bacterial pathogen and the site of infection. In the context of Mtb infections, type I IFN are considered to be detrimental to the host, and numerous recent studies have worked toward better understanding why. Surprisingly, we discovered an anti-microbial effect of type I IFN during Mtb infection in macrophages via the production of nitric oxide (NO). The role of NO in host resistance to tuberculosis has been extensively investigated and its bactericidal activity is attributed to the reactive nitrogen intermediates (RNI) such as NO2-, NO3- and peroxynitrite (ONOO-) (43, 44). Activation of macrophages with IFN-**γ** was especially powerful in augmenting RNI-mediated killing of Mtb (45-47). Two recent studies demonstrate that the major host protective role of NO during an in vivo infection with Mtb may not be its bactericidal activity but its immunosuppressive activity leading to reduced host tissue pathology (48, 49). NO is generated in the cell cytosol by NOS2 and it diffuses rapidly and freely at an estimated 5-10 cell lengths per second (50). This means that during an *in vivo* infection dilution is an important factor as within 1s the NO concentration has been diluted over 200 times in the NO-generating cell (50). In addition, the proteasome of Mtb has a role in resistance of the bacteria to RNI stress (51). It is thus possible that NO levels in the infected cells that actually generate the NO fail to accumulate to bactericidal threshold levels *in vivo* due to diffusion and dilution as compared to *ex vivo* infection experiments which are in a closed system and contain mostly infected cells. In any case, the potential of Mtb to inhibit IFN-**β**-mediated NO production will be advantageous for the pathogen. We do not believe that the recently reported direct bactericidal effect of IFN-**β** (52) plays a role in our observed bactericidal effects since otherwise the *Nos2*^-/-^ cells should not be different from wild-type BMDM (Figure 1E).

It seems confounding that Mtb would want to both induce IFN-**β** production and simultaneously inhibit it’s signaling. Considering that Mtb is a facultative intracellular pathogen, we hypothesize that inhibiting type I interferon signaling allows Mtb to mitigate the anti-bacterial effects of IFN-**β**-induced autocrine signaling (Figure 1). The transcriptome analysis of IFN-**β** regulated genes and their deregulation by Mtb infection suggests that there are some type I IFN responses that are detrimental to the bacterium during infection. CCL12/MCP-5 is one of the IFN-**β**-regulated cytokines that is strongly inhibited by Mtb infection at the mRNA and protein level (Figure 2B, 2E). It is a chemoattractant for monocytes and an agonist of CCR2 (53). Its expression on macrophages is induced by LPS and IFN-**γ** (53). Interestingly, another chemoattractant for monocytes, CCL5, is also upregulated via IFN-**β**-signaling and its expression is inhibited by Mtb (Figure S3). Consequently, the repression of these chemokines could lead to a reduction of monocyte recruitment within the Mtb-infected lungs. The type I IFN-driven expression of chemokines has been shown to be an important signal for recruitment of bone marrow monocytes to the site of infection with Listeria monocytogenes (54). Among other top down-regulated genes for which we also have confirmed reduced protein levels are the immunity-related GTPases Ifi204, Ifit1 and Iigp1 (Figure 2) (55). IIGP1 is involved in host defense against Chlamydia trachomatis and Toxoplasma gondii (56, 57). It is unlikely that this protein, at least in vivo, is involved in host defense against Mtb since *Iigp1*^-/-^ deficient mice do not have a phenotype (58). IFI204 is a cytosolic DNA sensor that binds to extracellular Mtb DNA and induces the cytosolic surveillance pathway via STING/TBK1/IRF3 signaling (17). Consequently, reduction of IFI204 expression by Mtb will reduce the activation of IRF3 and its regulon which includes *Ifn-**β*** transcription. This could thus represent an additional mechanism, in addition to the expression of the phosphodiesterase CdnP (23), by which Mtb regulates IFN-**β** production. Interestingly, *M. bovis* does not inhibit IFI204 expression and hence this host cell protein has an important role in host IFN-**β** production for this mycobacterial species (59). It is important to highlight, though, that the main cytosolic DNA sensor involved in recognition of Mtb DNA is cGAS (19-21). IFIT1 has well established anti-viral activity and is most strongly induced by IFN-**β** (60). Its activity is dependent upon selective binding of 5’-terminal regions of cap0-, cap1- and 5’ppp- mRNAs (61) and, hence, IFIT1 is an unlikely candidate for functioning in host cell resistance against Mtb infection. The expression of the *Nos2* gene is not affected by Mtb which we think is due to the early timepoint (4hpi) selected for the RNAseq analysis. *Nos2* can be induced by signaling via many TLRs and cytokine receptors (43, 62). In our system, Mtb infection causes a strong induction of TNF secretion which is amplified by addition of IFN-**β** (Figure S3). Consequently, we believe that TNF is the most likely cause for the observed late induction of NOS2, especially since it has been noted before that IFN-**β** and TNF synergistically mediate the induction of *Nos2* gene transcription (43, 62-64).

The capacity of Mtb to induce activation of STAT3 was recently reported (65) and the importance for STAT3 activation for virulence of Mtb was recently demonstrate by showing that deletion of STAT3 in myeloid cells increase resistance of mice to Mtb infection (66). STAT3 phosphorylation can inhibit type I IFN-mediated signaling, although this does not seem to occur at the level of STAT1/STAT2 phosphorylation but rather at the level of nuclear translocation (39, 40). Thus, STAT3 activation is likely not involved in our observed inhibition of TYK2/JAK1 phosphorylation by Mtb. However, we cannot exclude the possibility that Mtb exerts multiple strategies that synergize to prevent transcription of IFN-**β**-regulated genes and that STAT3-mediated inhibition may play a role during later time points to sustain the initial signaling inhibition. The capacity of STAT3 to inhibit *Nos2* gene expression may explain why NOS2 is only detected after 72h to 96h (Figure 1E). SOCS1, SOCS3, and USP18 are negative regulators of the IFNAR signaling which are typically induced at a later time point during IFN-**β** stimulation and serve as negative feedback regulators (3). Considering that our observed inhibition occurs already as early as 5 min post-infection, albeit an infection period of 4h, it is highly unlikely that the molecular mechanism of at least the early inhibition is dependent on these common negative regulators. It is known that Mtb possesses several phosphatases that have been shown to affect the host immune response by interfering with several signal transduction pathways. SapM is a phosphoinositide phosphatase that is essential in arresting phagosomal maturation by inhibiting phosphatidylinositol 3-phosphate phosphorylation (67). The tyrosine phosphatase PtpA is also involved inhibiting phagosomal maturation through inhibition of V-ATPase, and PtpB has been shown to inhibit ERK 1/2 and p38 signaling cascades (67-69). Although beyond the scope of this study, it would be interesting to investigate whether these proteins also regulate IFN-**β**-mediated signaling by directly dephosphorylating TYK2 or JAK1.

We here show that Mtb is susceptible to host cell IFNAR-signaling and has evolved to suppress it. In particular, Mtb inhibits IFNAR-mediated signaling at the level of the receptor-associated tyrosine kinases JAK1 and TYK2 (Figure 3). Nevertheless, this inhibition can be overcome by high concentrations of extracellular IFN-**β**, leading to reduced viability of intracellular Mtb. It is well established that Mtb induces the STING/TBK1/IRF3 signaling axis via extracellular Mtb DNA (eDNA) (17, 19-21), damage to host cell mitochondria leading to increase in cytosolic mitochondrial DNA (mtDNA) (24) or secretion of cyclic-di-AMP (c-di-AMP) (22, 23) (Figure 6). The capacity to produce IFN-**β** is associated with *in vivo* virulence of Mtb infections in mouse models and human studies (7, 12, 13, 15, 16). The levels of IFN-**β** production by mycobacteria-infected cells vary depending on the mycobacterial species and, in the context of Mtb, the specific strain that is used for the infection (24, 41, 42). High levels of IFN-**β** drive production of IL-10 and IL-1Ra (Figure 6) which antagonize the host protective activity of IL-1β and ultimately lead to increased host tissue destruction which establishes a replicative niche for Mtb (8). In contrast to the current dogma that Mtb induces production of type I IFNs in order to support its virulence, multiple studies have provided evidence that in some settings type I IFNs may have detrimental effects on Mtb. For example, *Isg15* is one of the most highly upregulated genes after type I IFNs stimulation of cells and it has a host protective role during Mtb infection (70). Furthermore, in the absence of IFN-**γ**, type I IFNs can promote the activation of macrophages for improved innate host response to Mtb (34). The strongest evidence for a host protective element in the IFN-**β** response was produced by showing that *Ifngr*^-/-^ *Ifnar*^-/-^ double-knock out mice are more susceptible when compared to *Ifn-*γ^-/-^ mice (6, 34, 71). IFN-**β** is protective during mouse infections against two non-tuberculous mycobacterial species (*M. smegmatis* and *M. avium ssp. Paratuberculosis*) (72). In this context, it is less surprising that Mtb has also evolved mechanisms to limit production of type I IFNs, possibly to achieve the “Goldilocks principle”. Overall, Mtb infection causes less production of IFN-**β** in *ex vivo* infected macrophages when compared to non-tuberculous mycobacteria (41). Indeed, we have shown previously that Mtb can inhibit *M. smegmatis-*induced IFN-**β** production in an ESX-1-dependent manner in bone marrow-derived dendritic cells (BMDCs) (41). There are potentially several mechanisms by which this inhibition occurs. The Mtb-mediated stimulation of TLR2 inhibits the induction of type I IFNs via TLR7/9 activation (73). In addition, the Mtb phosphodiesterase CdnP can reduce type I IFNs production by hydrolyzing bacterial-derived c-di-AMP and host-derived cGAMP, thereby limiting activation of the STING pathway (23). Importantly, this inhibition is relevant for full virulence of Mtb since a *CdnP* transposon mutant of Mtb is attenuated in mice (23).

**Fig 6.**
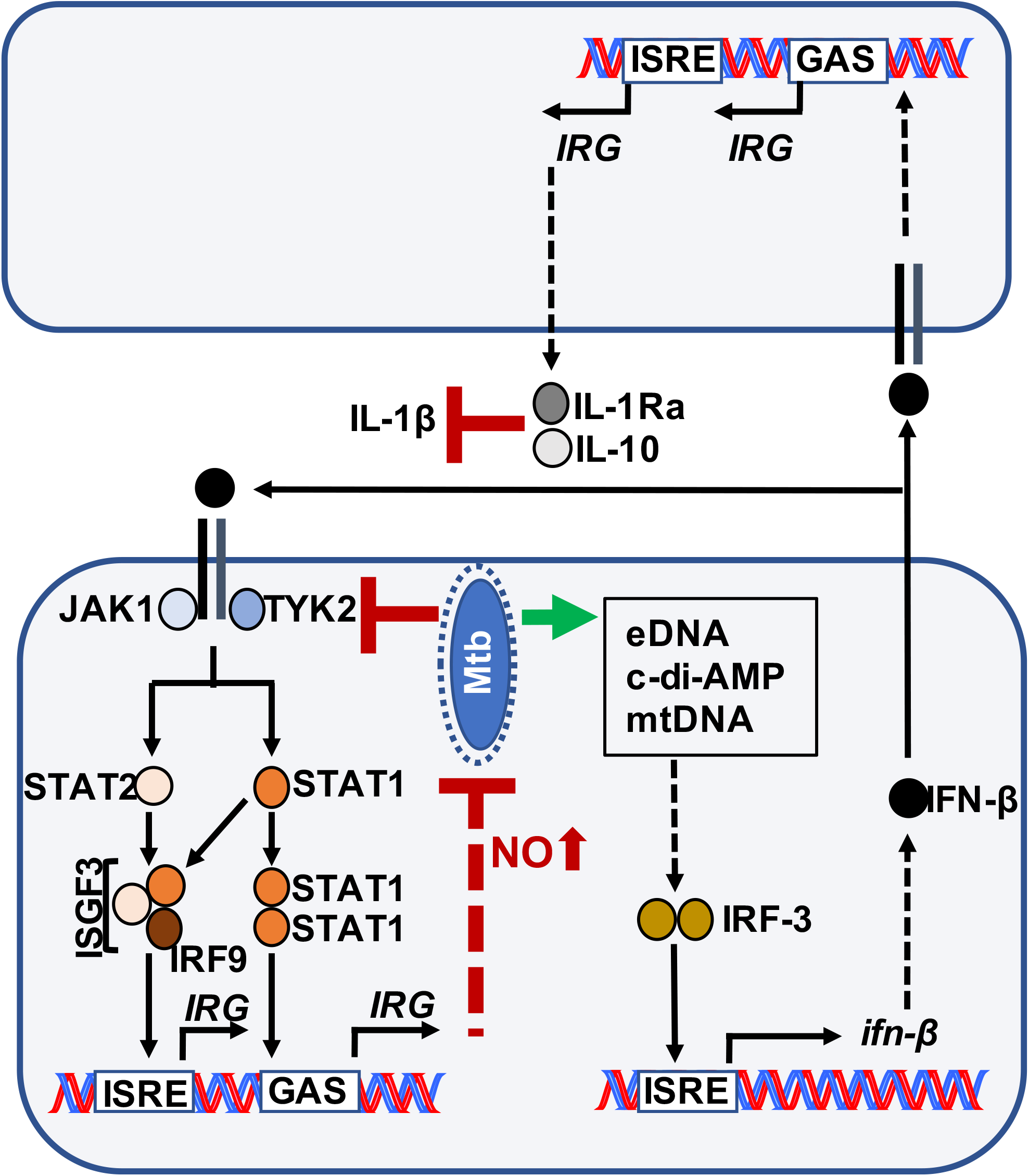
Overview of type I IFN signaling and production in Mtb-infected cells. Mtb induces the production of IFN-**β** in infected cells via release of bacterial DNA (eDNA) (17, 19-22) or damage to host cell mitochondria followed by increased mitochondrial DNA (mtDNA) in the cytosol (24). All these factors initiate the STING/TBK1 signaling pathway that leads to activation of the transcription factor IRF-3 and increased IFN-**β** production. The secreted IFN-**β** acts on bystander cells to increase secretion of IL-10 and IL-1Ra which lead to increased host cell necrosis and tissue damage, thus exacerbating the disease outcome (8). Here we show that Mtb inhibits autocrine IFNAR-signaling which limits not only the production of IFN-**β** but also the expression of genes with host cell defense properties such as *Nos2*.

We propose that Mtb has evolved to inhibit autocrine IFN-**β** signaling and its host protective effects in order to still take advantage of the benefits of paracrine IFN-**β** signaling on uninfected bystander cells, which is not inhibited by the Mtb-infected cells (Figure 5). This model might explain why in some settings IFN-**β** may be beneficial for the host (6); for example, in the case of nontuberculous mycobacteria which cannot inhibit IFNAR-mediated signaling (e.g. *M. smegmatis*) and who are susceptible to a host type I IFN response (72).

## Acknowledgements

We would like to thank the UMD Genomics Core facility for excellent technical assistance. Dr. Sophie Helaine (Imperial College, London) for advice and editing of manuscript.

## References

1. Diamond, M. S., and M. Farzan. 2012. The broad-spectrum antiviral functions of IFIT and IFITM proteins. Nat. Rev. Immunol. 13: 46–57.

2. McNab, F., K. Mayer-Barber, A. Sher, A. Wack, and A. O’Garra. 2015. Type I interferons in infectious disease. Nat. Rev. Immunol. 15: 87–103.

3. Ivashkiv, L. B., and L. T. Donlin. 2014. Regulation of type I interferon responses. Nat. Rev. Immunol. 14: 36–49.

4. Boxx, G. M., and G. Cheng. 2016. The Roles of Type I Interferon in Bacterial Infection. Cell Host and Microbe 19: 760–769.

5. Kovarik, P., V. Castiglia, M. Ivin, and F. Ebner. 2016. Type I Interferons in Bacterial Infections: A Balancing Act. Front. Immunol. 7: 307–8.

6. Moreira-Teixeira, L., K. Mayer-Barber, A. Sher, and A. O’Garra. 2018. Type I interferons in tuberculosis: Foe and occasionally friend. J. Exp. Med. 215: 1273–1285.

7. Dorhoi, A., V. Yeremeev, G. Nouailles, J. Weiner, S. Jörg, E. Heinemann, D. Oberbeck-Müller, J. K. Knaul, A. Vogelzang, S. T. Reece, K. Hahnke, H.-J. Mollenkopf, V. Brinkmann, and S. H. E. Kaufmann. 2014. Type I IFN signaling triggers immunopathology in tuberculosis-susceptible mice by modulating lung phagocyte dynamics. Eur. J. Immunol. 44: 2380–2393.

8. Mayer-Barber, K. D., B. B. Andrade, S. D. Oland, E. P. Amaral, D. L. Barber, J. Gonzales, S.C. Derrick, R. Shi, N. P. Kumar, W. Wei, X. Yuan, G. Zhang, Y. Cai, S. Babu, M. Catalfamo, A. M. Salazar, L. E. Via, C. E. Barry, and A. Sher. 2014. Host-directed therapy of tuberculosis based on interleukin-1 and type I interferon crosstalk. Nature 511: 99–103.

9. Mayer-Barber, K. D., B. B. Andrade, D. L. Barber, S. Hieny, C. G. Feng, P. Caspar, S. Oland, S. Gordon, and A. Sher. 2011. Innate and adaptive interferons suppress IL-1α and IL-1β production by distinct pulmonary myeloid subsets during Mycobacterium tuberculosis infection. Immunity 35: 1023–1034.

10. Novikov, A., M. Cardone, R. Thompson, K. Shenderov, K. D. Kirschman, K. D. Mayer-Barber, T. G. Myers, R. L. Rabin, G. Trinchieri, A. Sher, and C. G. Feng. 2011. Mycobacterium tuberculosis triggers host type I IFN signaling to regulate IL-1β production in human macrophages. J. Immunol. 187: 2540–2547.

11. Maertzdorf, J., D. Repsilber, S. K. Parida, K. Stanley, T. Roberts, G. Black, G. Walzl, and S. H. E. Kaufmann. 2010. Human gene expression profiles of susceptibility and resistance in tuberculosis. Genes Immun 12: 15–22.

12. Berry, M. P. R., C. M. Graham, F. W. McNab, Z. Xu, S. A. A. Bloch, T. Oni, K. A. Wilkinson,R. Banchereau, J. Skinner, R. J. Wilkinson, C. Quinn, D. Blankenship, R. Dhawan, J. J. Cush,A. Mejias, O. Ramilo, O. M. Kon, V. Pascual, J. Banchereau, D. Chaussabel, and A. O’Garra. 2010. An interferon-inducible neutrophil-driven blood transcriptional signature in human tuberculosis. Nature 466: 973–977.

13. Antonelli, L. R. V., A. Gigliotti Rothfuchs, R. Gonçalves, E. Roffê, A. W. Cheever, A. Bafica,A. M. Salazar, C. G. Feng, and A. Sher. 2010. Intranasal Poly-IC treatment exacerbates tuberculosis in mice through the pulmonary recruitment of a pathogen-permissive monocyte/macrophage population. J. Clin. Invest. 120: 1674–1682.

14. Ordway, D., M. Henao-Tamayo, M. Harton, G. Palanisamy, J. Troudt, C. Shanley, R. J. Basaraba, and I. M. Orme. 2007. The hypervirulent Mycobacterium tuberculosis strain HN878 induces a potent TH1 response followed by rapid down-regulation. J. Immunol. 179: 522–531.

15. Stanley, S. A., J. E. Johndrow, P. Manzanillo, and J. S. Cox. 2007. The Type I IFN response to infection with Mycobacterium tuberculosis requires ESX-1-mediated secretion and contributes to pathogenesis. J. Immunol. 178: 3143–3152.

16. Manca, C., L. Tsenova, A. Bergtold, S. Freeman, M. Tovey, J. M. Musser, C. E. Barry, V.H. Freedman, and G. Kaplan. 2001. Virulence of a Mycobacterium tuberculosis clinical isolate in mice is determined by failure to induce Th1 type immunity and is associated with induction of IFN-alpha/beta. Proc. Natl. Acad. Sci. U.S.A. 98: 5752–5757.

17. Manzanillo, P. S., M. U. Shiloh, D. A. Portnoy, and J. S. Cox. 2012. Mycobacterium tuberculosis activates the DNA-dependent cytosolic surveillance pathway within macrophages. Cell Host and Microbe 11: 469–480.

18. Watson, R. O., P. S. Manzanillo, and J. S. Cox. 2012. Extracellular M. tuberculosis DNA targets bacteria for autophagy by activating the host DNA-sensing pathway. Cell 150: 803–815.

19. Wassermann, R., M. F. Gulen, C. Sala, S. G. Perin, Y. Lou, J. Rybniker, J. L. Schmid-Burgk, T. Schmidt, V. Hornung, S. T. Cole, and A. Ablasser. 2015. Mycobacterium tuberculosis Differentially Activates cGAS- and Inflammasome-Dependent Intracellular Immune Responses through ESX-1. Cell Host and Microbe 17: 799–810.

20. Watson, R. O., S. L. Bell, D. A. MacDuff, J. M. Kimmey, E. J. Diner, J. Olivas, R. E. Vance,C. L. Stallings, H. W. Virgin, and J. S. Cox. 2015. The Cytosolic Sensor cGAS Detects Mycobacterium tuberculosis DNA to Induce Type I Interferons and Activate Autophagy. Cell Host and Microbe 17: 811–819.

21. Collins, A. C., H. Cai, T. Li, L. H. Franco, X.-D. Li, V. R. Nair, C. R. Scharn, C. E. Stamm,B. Levine, Z. J. Chen, and M. U. Shiloh. 2015. Cyclic GMP-AMP Synthase Is an Innate Immune DNA Sensor for Mycobacterium tuberculosis. Cell Host and Microbe 17: 820–828.

22. Dey, B., R. J. Dey, L. S. Cheung, S. Pokkali, H. Guo, J.-H. Lee, and W. R. Bishai. 2015. A bacterial cyclic dinucleotide activates the cytosolic surveillance pathway and mediates innate resistance to tuberculosis. Nat. Med. 21: 401–406.

23. Dey, R. J., B. Dey, Y. Zheng, L. S. Cheung, J. Zhou, D. Sayre, P. Kumar, H. Guo, G. Lamichhane, H. O. Sintim, and W. R. Bishai. 2017. Inhibition of innate immune cytosolic surveillance by an M. tuberculosis phosphodiesterase. Nat Chem Biol 13: 210–217.

24. Wiens, K. E., and J. D. Ernst. 2016. The Mechanism for Type I Interferon Induction by Mycobacterium tuberculosis is Bacterial Strain-Dependent. PLoS Pathog 12: e1005809–20.

25. Deonarain, R., A. Alcami, M. Alexiou, M. J. Dallman, D. R. Gewert, and A. C. G. Porter. 2000. Impaired Antiviral Response and Alpha/Beta Interferon Induction in Mice Lacking Beta Interferon. Journal of Virology 74: 3404–3409.

26. Srinivasan, L., S. A. Gurses, B. E. Hurley, J. L. Miller, P. C. Karakousis, and V. Briken. 2016. Identification of a Transcription Factor That Regulates Host Cell Exit and Virulence of Mycobacterium tuberculosis. PLoS Pathog 12: e1005652.

27. Bolger, A. M., M. Lohse, and B. Usadel. 2014. Trimmomatic: a flexible trimmer for Illumina sequence data. Bioinformatics 30: 2114–2120.

28. Yates, A., W. Akanni, M. R. Amode, D. Barrell, K. Billis, D. Carvalho-Silva, C. Cummins, P. Clapham, S. Fitzgerald, L. Gil, C. G. Girón, L. Gordon, T. Hourlier, S. E. Hunt, S. H. Janacek,N. Johnson, T. Juettemann, S. Keenan, I. Lavidas, F. J. Martin, T. Maurel, W. McLaren, D. N. Murphy, R. Nag, M. Nuhn, A. Parker, M. Patricio, M. Pignatelli, M. Rahtz, H. S. Riat, D. Sheppard, K. Taylor, A. Thormann, A. Vullo, S. P. Wilder, A. Zadissa, E. Birney, J. Harrow, M. Muffato, E. Perry, M. Ruffier, G. Spudich, S. J. Trevanion, F. Cunningham, B. L. Aken, D. R. Zerbino, and P. Flicek. 2016. Ensembl 2016. Nucleic Acids Research 44: D710–D716.

29. Kim, D., G. Pertea, C. Trapnell, H. Pimentel, R. Kelley, and S. L. Salzberg. 2013. TopHat2: accurate alignment of transcriptomes in the presence of insertions, deletions and gene fusions. Genome Biol. 14: R36.

30. Anders, S., P. T. Pyl, and W. Huber. 2015. HTSeq–a Python framework to work with high-throughput sequencing data. Bioinformatics 31: 166–169.

31. Gentleman, R. C., V. J. Carey, D. M. Bates, B. Bolstad, M. Dettling, S. Dudoit, B. Ellis, L. Gautier, Y. Ge, J. Gentry, K. Hornik, T. Hothorn, W. Huber, S. Iacus, R. Irizarry, F. Leisch, C. Li, M. Maechler, A. J. Rossini, G. Sawitzki, C. Smith, G. Smyth, L. Tierney, J. Y. H. Yang, and J. Zhang. 2004. Bioconductor: open software development for computational biology and bioinformatics. Genome Biol. 5: R80.

32. Law, C. W., Y. Chen, W. Shi, and G. K. Smyth. 2014. voom: Precision weights unlock linear model analysis tools for RNA-seq read counts. Genome Biol. 15: R29.

33. Leek, J. T., W. E. Johnson, H. S. Parker, A. E. Jaffe, and J. D. Storey. 2012. The sva package for removing batch effects and other unwanted variation in high-throughput experiments. Bioinformatics 28: 882–883.

34. Moreira-Teixeira, L., J. Sousa, F. W. McNab, E. Torrado, F. Cardoso, H. Machado, F. Castro, V. Cardoso, J. Gaifem, X. Wu, R. Appelberg, A. G. Castro, A. O’Garra, and M. Saraiva. 2016. Type I IFN Inhibits Alternative Macrophage Activation during Mycobacterium tuberculosis Infection and Leads to Enhanced Protection in the Absence of IFN-γ Signaling. J. Immunol. 197: 4714–4726.

35. Evans, J. D., R. A. Crown, J. A. Sohn, and C. Seeger. 2011. West Nile virus infection induces depletion of IFNAR1 protein levels. Viral Immunol. 24: 253–263.

36. Xia, C., M. Vijayan, C. J. Pritzl, S. Y. Fuchs, A. B. McDermott, and B. Hahm. 2015. Hemagglutinin of Influenza A Virus Antagonizes Type I Interferon (IFN) Responses by Inducing Degradation of Type I IFN Receptor 1. Journal of Virology 90: 2403–2417.

37. Ting, L. M., A. C. Kim, A. Cattamanchi, and J. D. Ernst. 1999. Mycobacterium tuberculosis inhibits IFN-gamma transcriptional responses without inhibiting activation of STAT1. J. Immunol. 163: 3898–3906.

38. Villarino, A. V., Y. Kanno, and J. J. O’Shea. 2017. Mechanisms and consequences of Jak–STAT signaling in the immune system. Nat Immunol 18: 374–384.

39. Ho, H. H., and L. B. Ivashkiv. 2006. Role of STAT3 in Type I Interferon Responses. J. Biol. Chem. 281: 14111–14118.

40. Wang, W.-B., Wang, W. B., D. E. Levy, D. E. Levy, C. K. Lee, and C.-K. Lee. 2011. STAT3 Negatively Regulates Type I IFN-Mediated Antiviral Response. J. Immunol. 187: 2578–2585.

41. Shah, S., A. Bohsali, S. E. Ahlbrand, L. Srinivasan, V. A. K. Rathinam, S. N. Vogel, K. A. Fitzgerald, F. S. Sutterwala, and V. Briken. 2013. Cutting edge: Mycobacterium tuberculosis but not nonvirulent mycobacteria inhibits IFN-β and AIM2 inflammasome-dependent IL-1β production via its ESX-1 secretion system. J. Immunol. 191: 3514–3518.

42. Manca, C., L. Tsenova, S. Freeman, A. K. Barczak, M. Tovey, P. J. Murray, C. Barry III, and G. Kaplan. 2005. Hypervirulent M. tuberculosis W/Beijing strains upregulate type I IFNs and increase expression of negative regulators of the Jak-Stat pathway. J. Interferon Cytokine Res. 25: 694–701.

43. MacMicking, J., Q. W. Xie, and C. Nathan. 1997. Nitric oxide and macrophage function. Annu. Rev. Immunol. 15: 323–350.

44. Jamaati, H., E. Mortaz, Z. Pajouhi, G. Folkerts, M. Movassaghi, M. Moloudizargari, I. M. Adcock, and J. Garssen. 2017. Nitric Oxide in the Pathogenesis and Treatment of Tuberculosis. Front Microbiol 8: 436–11.

45. Flesch, I. E., and S. H. Kaufmann. 1991. Mechanisms involved in mycobacterial growth inhibition by gamma interferon-activated bone marrow macrophages: role of reactive nitrogen intermediates. Infection and Immunity 59: 3213–3218.

46. Denis, M. 1991. Interferon-gamma-treated murine macrophages inhibit growth of tubercle bacilli via the generation of reactive nitrogen intermediates. Cell. Immunol. 132: 150–157.

47. Chan, J., Y. Xing, R. S. Magliozzo, and B. R. Bloom. 1992. Killing of virulent Mycobacterium tuberculosis by reactive nitrogen intermediates produced by activated murine macrophages. J. Exp. Med. 175: 1111–1122.

48. Mishra, B. B., R. R. Lovewell, A. J. Olive, G. Zhang, W. Wang, E. Eugenin, C. M. Smith, J.Y. Phuah, J. E. Long, M. L. Dubuke, S. G. Palace, J. D. Goguen, R. E. Baker, S. Nambi, R. Mishra, M. G. Booty, C. E. Baer, S. A. Shaffer, V. Dartois, B. A. McCormick, X. Chen, and C.M. Sassetti. 2017. Nitric oxide prevents a pathogen-permissive granulocytic inflammation during tuberculosis. Nature Microbiology 1–11.

49. Mishra, B. B., V. A. K. Rathinam, G. W. Martens, A. J. Martinot, H. Kornfeld, K. A. Fitzgerald, and C. M. Sassetti. 2012. Nitric oxide controls the immunopathology of tuberculosis by inhibiting NLRP3 inflammasome–dependent processing of IL-1β. Nat Immunol 14: 52–60.

50. Thomas, D. D., L. A. Ridnour, J. S. Isenberg, W. Flores-Santana, C. H. Switzer, S. Donzelli, P. Hussain, C. Vecoli, N. Paolocci, S. Ambs, C. A. Colton, C. C. Harris, D. D. Roberts, and D. A. Wink. 2008. The chemical biology of nitric oxide: implications in cellular signaling. Free Radical Biology and Medicine 45: 18–31.

51. Darwin, K. H., S. Ehrt, J.-C. Gutierrez-Ramos, N. Weich, and C. F. Nathan. 2003. The proteasome of Mycobacterium tuberculosis is required for resistance to nitric oxide. Science 302: 1963–1966.

52. Kaplan, A., M. W. Lee, A. J. Wolf, J. J. Limon, C. A. Becker, M. Ding, R. Murali, E. Y. Lee,G. Y. Liu, G. C. L. Wong, and D. M. Underhill. 2017. Direct Antimicrobial Activity of IFN-β. J. Immunol. 198: 4036–4045.

53. Sarafi, M. N., E. A. Garcia-Zepeda, J. A. MacLean, I. F. Charo, and A. D. Luster. 1997. Murine monocyte chemoattractant protein (MCP)-5: a novel CC chemokine that is a structural and functional homologue of human MCP-1. J. Exp. Med. 185: 99–109.

54. Jia, T., I. Leiner, G. Dorothee, K. Brandl, and E. G. Pamer. 2009. MyD88 and Type I interferon receptor-mediated chemokine induction and monocyte recruitment during Listeria monocytogenes infection. J. Immunol. 183: 1271–1278.

55. Kim, B.-H., J. D. Chee, C. J. Bradfield, E.-S. Park, P. Kumar, and J. D. MacMicking. 2016. Interferon-induced guanylate-binding proteins in inflammasome activation and host defense. Nat Immunol 17: 481–489.

56. Martens, S., I. Parvanova, J. Zerrahn, G. Griffiths, G. Schell, G. Reichmann, and J. C. Howard. 2005. Disruption of Toxoplasma gondii Parasitophorous Vacuoles by the Mouse p47-Resistance GTPases. PLoS Pathog 1: e24–15.

57. Al-Zeer, M. A., H. M. Al-Younes, P. R. Braun, J. Zerrahn, and T. F. Meyer. 2009. IFN-γ-Inducible Irga6 Mediates Host Resistance against Chlamydia trachomatis via Autophagy. PLoS ONE 4: e4588–13.

58. Liesenfeld, O., I. Parvanova, J. Zerrahn, S.-J. Han, F. Heinrich, M. Muñoz, F. Kaiser, T. Aebischer, T. Buch, A. Waisman, G. Reichmann, O. Utermöhlen, E. von Stebut, F. D. von Loewenich, C. Bogdan, S. Specht, M. Saeftel, A. Hoerauf, M. M. Mota, S. Könen-Waisman, S. H. E. Kaufmann, and J. C. Howard. 2011. The IFN-γ-Inducible GTPase, Irga6, Protects Mice against Toxoplasma gondii but Not against Plasmodium berghei and Some Other Intracellular Pathogens. PLoS ONE 6: e20568–12.

59. Chunfa, L., S. Xin, L. Qiang, S. Sreevatsan, L. Yang, D. Zhao, and X. Zhou. 2017. The Central Role of IFI204 in IFN-β Release and Autophagy Activation during Mycobacterium bovis Infection. Front Cell Infect Microbiol 7: 169.

60. Fensterl, V., and G. C. Sen. 2015. Interferon-induced Ifit proteins: their role in viral pathogenesis. Journal of Virology 89: 2462–2468.

61. Kumar, P., T. R. Sweeney, M. A. Skabkin, O. V. Skabkina, C. U. T. Hellen, and T. V. Pestova. 2014. Inhibition of translation by IFIT family members is determined by their ability to interact selectively with the 5“-terminal regions of cap0-, cap1- and 5”ppp- mRNAs. Nucleic Acids Research 42: 3228–3245.

62. Bogdan, C. 2001. Nitric oxide and the immune response. Nat Immunol 2: 907–916.

63. Farlik, M., B. Reutterer, C. Schindler, F. Greten, C. Vogl, M. Müller, and T. Decker. 2010. Nonconventional initiation complex assembly by STAT and NF-kappaB transcription factors regulates nitric oxide synthase expression. Immunity 33: 25–34.

64. Bachmann, M., Z. Waibler, T. Pleli, J. Pfeilschifter, and H. Mühl. 2017. Type I Interferon Supports Inducible Nitric Oxide Synthase in Murine Hepatoma Cells and Hepatocytes and during Experimental Acetaminophen-Induced Liver Damage. Front. Immunol. 8: 75–11.

65. Queval, C. J., O.-R. Song, N. Deboosère, V. Delorme, A.-S. Debrie, R. Iantomasi, R. Veyron-Churlet, S. Jouny, K. Redhage, G. Deloison, A. Baulard, M. Chamaillard, C. Locht, and P. Brodin. 2016. STAT3 Represses Nitric Oxide Synthesis in Human Macrophages upon Mycobacterium tuberculosis Infection. Sci. Rep. 1–14.

66. Gao, Y., J. I. Basile, C. Classon, D. Gavier-Widen, A. Yoshimura, B. Carow, and M. E. Rottenberg. 2018. STAT3 expression by myeloid cells is detrimental for the T-cell-mediated control of infection with Mycobacterium tuberculosis. PLoS Pathog 14: e1006809.

67. Wong, D., J. D. Chao, and Y. Av-Gay. 2013. Mycobacterium tuberculosis-secreted phosphatases: from pathogenesis to targets for TB drug development. Trends in microbiology 21: 100–109.

68. Zhou, B., Y. He, X. Zhang, J. Xu, Y. Luo, Y. Wang, S. G. Franzblau, Z. Yang, R. J. Chan, Y. Liu, J. Zheng, and Z.-Y. Zhang. 2010. Targeting mycobacterium protein tyrosine phosphatase B for antituberculosis agents. Proceedings of the National Academy of Sciences 107: 4573–4578.

69. Wong, D., H. Bach, J. Sun, Z. Hmama, and Y. Av-Gay. 2011. Mycobacterium tuberculosis protein tyrosine phosphatase (PtpA) excludes host vacuolar-H+-ATPase to inhibit phagosome acidification. Proceedings of the National Academy of Sciences 108: 19371–19376.

70. Kimmey, J. M., J. A. Campbell, L. A. Weiss, K. J. Monte, D. J. Lenschow, and C. L. Stallings. 2017. The impact of ISGylation during Mycobacterium tuberculosis infection in mice. Microbes Infect.

71. Desvignes, L., A. J. Wolf, and J. D. Ernst. 2012. Dynamic roles of type I and type II IFNs in early infection with Mycobacterium tuberculosis. J. Immunol. 188: 6205–6215.

72. Ruangkiattikul, N., A. Nerlich, K. Abdissa, S. Lienenklaus, A. Suwandi, N. Janze, K. Laarmann, J. Spanier, U. Kalinke, S. Weiss, and R. Goethe. 2017. cGAS-STING-TBK1-IRF3/7 induced interferon-β contributes to the clearing of non tuberculous mycobacterial infection in mice. Virulence 8: 1–13.

73. Liu, Y. C., D. P. Simmons, X. Li, D. W. Abbott, W. H. Boom, and C. V. Harding. 2012. TLR2 signaling depletes IRAK1 and inhibits induction of type I IFN by TLR7/9. J. Immunol. 188: 1019–1026.

